# Three-dimensional observations of an aperiodic oscillatory gliding motility behaviour in *Myxococcus xanthus* using confocal interference reflection microscopy

**DOI:** 10.1101/722231

**Authors:** Liam M. Rooney, Lisa S. Kölln, Ross Scrimgeour, William B. Amos, Paul A. Hoskisson, Gail McConnell

## Abstract

The Delta-proteobacterium, *Myxococcus xanthus*, has been used as a model for bacterial motility and to provide insights of bacterial swarming behaviours. Fluorescence microscopy techniques have shown that various mechanisms are involved in gliding motility, but these have almost entirely been limited to 2D studies and there is currently no understanding of gliding motility in a 3D context. We present here the first use of confocal interference reflection microscopy (IRM) to study gliding bacteria, and we reveal aperiodic oscillatory behaviour with changes in the position of the basal membrane relative to the coverglass on the order of 90 nm *in vitro*. Firstly, we use a model plano-convex lens specimen to show how topological information can be obtained from the wavelength-dependent interference pattern in IRM. We then use IRM to observe gliding *M. xanthus* and show that cells undergo previously unobserved changes in their height as they glide. We compare the wild-type with mutants of reduced motility, which also exhibit the same changes in adhesion profile during gliding. We find that the general gliding behaviour is independent of the proton motive force-generating complex, AglRQS, and suggest that the novel behaviour we present here may be a result of recoil and force transmission along the length of the cell body following firing of the Type IV pili.

## Introduction

Bacteria use a number of mechanisms to move through their local environment in response to chemotactic signals, to form communities or to invade their host. The most studied mode of bacterial motility is flagellar-mediated movement. Other modes, such as the twitching motility displayed by *Pseudomonas aeruginosa*, use Type IV pili (T4P) to direct movement based on the extension, adhesion and retraction of polar filaments from the leading pole of the cell^1,2^. However, not all bacteria rely solely on extracellular appendages for motility. The phenomenon of gliding motility has been identified in a diverse range of bacterial species spanning various phyla^3–8^. The Delta-proteobacterium *Myxococcus xanthus* displays two different modes of gliding motility - adventurous motility and social motility - to seek out nutrients or prey as part of its complex lifecycle^4,9–15^.

There are contrasting models proposed to explain the mechanisms underpinning gliding motility^9,16–19^. The focal adhesion complex (FAC) model proposes that FACs form on the basal surface of the cell and attach to the underlying substrate while coupling to the helical MreB cytoskeleton on the cell’s inner-membrane^16,17,20,21^. It has been shown that FACs translocate linearly from the leading pole as the cell moves forwards and is driven by the force generated by the AglRQS gliding complex, which is associated with the FAC^16,22^. The FAC model requires the basal layer of the cell to be firmly attached to the underlying substrate, however, it remains unclear how the complex is able to traverse the peptidoglycan cell wall without compromising the structural integrity of the cell^8,19^. A second model has been suggested where proton motive force (PMF) generated by AglRQS results in a helical rotation of the MreB cytoskeleton in gliding cells which are firmly adhered to a solid substrate^9,22–25^. In the helical rotation model, stationary foci of fluorescently-tagged motor complex subunits have been explained as being a build-up of multiple complexes arrayed in “traffic jams” which result from areas of differing resistance in the underlying substrate ^9,24^. Both models converge where the gliding cell is adhered firmly to the surface of the underlying substrate to facilitate gliding. However, our observations show that cells are not in fact firmly adhered during gliding motility, but instead exhibit aperiodic fluctuations in their height as they glide.

Bacterial gliding motility has mainly been studied using phase contrast and fluorescence microscopy techniques which do not provide 3D information about cell movement^9,17,22,26^. We hypothesised that axial changes in cell shape during gliding motility may occur due to the complex nature of underlying mechanisms such as FAC translocation and bulk movement of the cytoskeleton. We reasoned that novel 3D behaviours could be visualised using the label-free microscopy technique interference reflection microscopy (IRM). This technique has previously been used to study focal adhesion sites of eukaryotic cells on glass substrates^27–34^ and microtubule dynamics^35–37^. Previous studies have used widefield IRM to observe gliding motility in *Cytophaga spp.*, where rotation and adhesion to glass surfaces were characterised^4^. Twitching motility in *Pseudomonas fluorescens* has also been investigated using widefield IRM, where the attachment profile of twitching cells was found to be dependent on the presence of different electrolytes^38^. However, these previous widefield IRM studies have a low contrast between the orders of interference, arising from the short coherence length of the light source. The contrast of higher order interference fringes can be improved by using IRM in confocal mode where coherent laser light is used and out of focus signal is significantly reduced by incorporating a pinhole before the photodetector ^39^.

In IRM, the detected signal originates from the interference of reflected light at refractive index boundaries within a live cell specimen that is plated on a glass coverslip. Constructive and destructive interference of reflected light originating from the coverslip-medium and medium-cell interfaces result in a fringe pattern which can be translated to the axial position of the cell surface^40,41^. Using IRM, a five-fold axial resolution enhancement is achieved over conventional widefield or point-scanning microscopy techniques, where 3D information can be extracted from a 2D image thus overcoming the limitations in optical sectioning of thin specimens (i.e. bacteria)^27,40,41^. An additional benefit of IRM in comparison to other similar methods is that it requires almost no adaptation to a standard confocal or widefield microscope and is compatible with existing objective lenses.

The application of IRM to biological specimens has been documented since the 1960s, but the interpretation of IRM data can be difficult^27^. Theoretically, the axial resolution of IRM can be as high as 15 nm^27^, but in practice factors such as dense protein aggregates, transport of dry mass to the basal membrane, changes in local membrane density and proximity of intracellular structures to the membrane affect the brightness of the detected IRM image^40,42^. However, in thin specimens such as bacterial cells, where internal shifting of dry mass is unlikely due to the lack of intracellular vesicular transport, IRM remains a viable height-measuring technique^43^. Godwin *et al.* found that separation between the cell and the coverslip on the order of 100 nm can be easily distinguished without the influence from ambiguities introduced by above listed factors^4^. Others have disputed the capability of IRM to accurately measure close contact sites in live cells, for example, by imaging the displacement of a thin layer of fluorescent dye between adherent cells and the coverglass with TIRF microscopy^44^. In the present work it is assumed that the IRM contrast of bacterial specimens is an indication of height of the cell surface above the substrate.

This study is the first application of confocal IRM to bacterial specimens. However, previous studies have used similar techniques as a means of contrast enhancement. One such study used reflection interference contrast microscopy (RICM), where polarising filters and an antiflex objective are used to filter out reflected light from outside the specimen plane^39,45,46^, to only image *M. xanthus* surface detachment from the coverglass^17^. Total internal reflection fluorescence (TIRF) microscopy has also shown that FACs which attach to and rotate the MreB cytoskeleton are found in distinct foci on the basal side of the cell^9,16^. These membrane-associated complexes have been suggested to change the surface topology of the gliding cell depending on the cargo load of the molecular motor^9^. More recently, interferometric scattering microscopy (iSCAT), which detects both the reflected and scattered light, has been used to observe T4P-mediated twitching motility in *P. aeruginosa*. In this work the authors generated 3D illustrations which revealed the role of T4P machinery subunits in extension, attachment and retraction based on the interference pattern in iSCAT images^47^.

## Background Theory

For the image formation theory in IRM, a simplified three-layer model system is assumed that only consists of the coverglass (*g*), the cell medium (*m*) and the cell (*c*). In this model, the cell medium can be viewed as a thin film with varying height, dependent on how closely the cell is attached to the coverslip. The intensity of the reflected light *I*_*g-m*_ at an interface, for example, the coverglass-medium interface (*g-m*), follows the Fresnel equations:

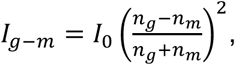

with *I*_*0*_ being the intensity of the incident light beam and *n*_*g/m*_ as the respective refractive indices of the adjacent materials.

The reflected light beams at the coverglass-medium (*g-m*) and the medium-cell (*m-c*) interface coincide, leading to de-/constructive interference dependent on the optical path length difference *z* between the two beams (Supplementary Figure 1). The intensity follows *I(z)*:

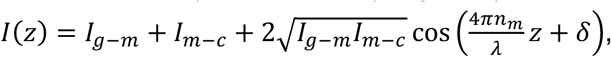

with *n*_*m*_ as the refractive index of the cell medium, *λ* as the wavelength of the light and *δ* as the phase difference^48^. Since the refractive index of the cell medium is smaller than the refractive index of the cell, a phase shift occurs upon reflection so that *δ* equals π^40^. Accordingly, deconstructive interference occurs at optical path length differences of 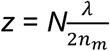 and constructive interference occurs at 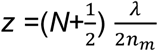with *N* = 0, 1, 2, … as the interference order. This wavelength-dependency can be used to extract the cell topology from IRM images since overlap of the interference fringes of different wavelengths decreases with increasing distance between the cell and the coverglass. The decreasing fringe overlap with increasing separation between the cell and coverglass, as observable in colour-merged IRM images obtained for different wavelengths, results in a clear colour-ordering along the cell body when the cell is lifted up from the coverglass.

Common assumptions of this IRM model are that no other refractive index boundary exists in the cell specimen (i.e. that the refractive index of the cell is constant) and that incident and reflected light are perpendicular to the coverglass^28^. Also, the impact of the numerical aperture (NA) is neglected which affects the depth of field that is imaged. In IRM, high NA objectives are used to limit detection of reflection signals to those originating from interfaces close to the coverglass, which establishes an experimental condition that is close to the three-layer system established, here, consisting of the coverglass, the cell medium and the cell body^39^.

## Results

### Characterisation of model specimen

We characterised the axial height intensity profile of a specimen of known structure to compare with IRM theory by acquiring IRM images at different wavelengths of a plano-convex lens (focal length = 72 mm) that was placed on a coverglass. A composite of IRM images of the lens specimen acquired at 488 nm, 514 nm and 543 nm is shown in Figure 1a. In Figure 1b, a cross-sectional schematic of a plano-convex lens specimen is shown, outlining the axial position of the intensity maxima that are caused by constructive interference for different order and wavelength. We analysed the intensity of the interference fringes by comparing the radial intensity profile with the theory of IRM regarding fringe separation^27^. We calculated that the theoretical spacing (λ/2*n*) between the intensity maxima caused by constructive interference for the different wavelengths are 244 nm, 258 nm and 272 nm (*n* = 1). With the experimental data we obtained slightly different spacings for the intensity maxima, with 249 ± 1 nm, 262 ± 1 nm and 277 ± 1 nm (Figure 2b). Thus, experimental and expected values deviated by 2.07%, 1.63% and 1.91% for the different wavelengths. Additionally, the overlap of the intensity maxima of different acquisition wavelengths decreased with lens-to-coverslip distance (Figure 2). These observations provided a sense of directionality regarding specimen topology, where the curvature of the plano-convex lens specimen was clear from the acquired images and allowed for 3D reconstruction of the lens specimen (Figure 2c). This means that assumptions outlined in Background Theory are appropriate to reconstruct the morphology of simple model systems. Here a multi-wavelength IRM approach provides important additional morphological information over single wavelength IRM.

**Figure 1.**
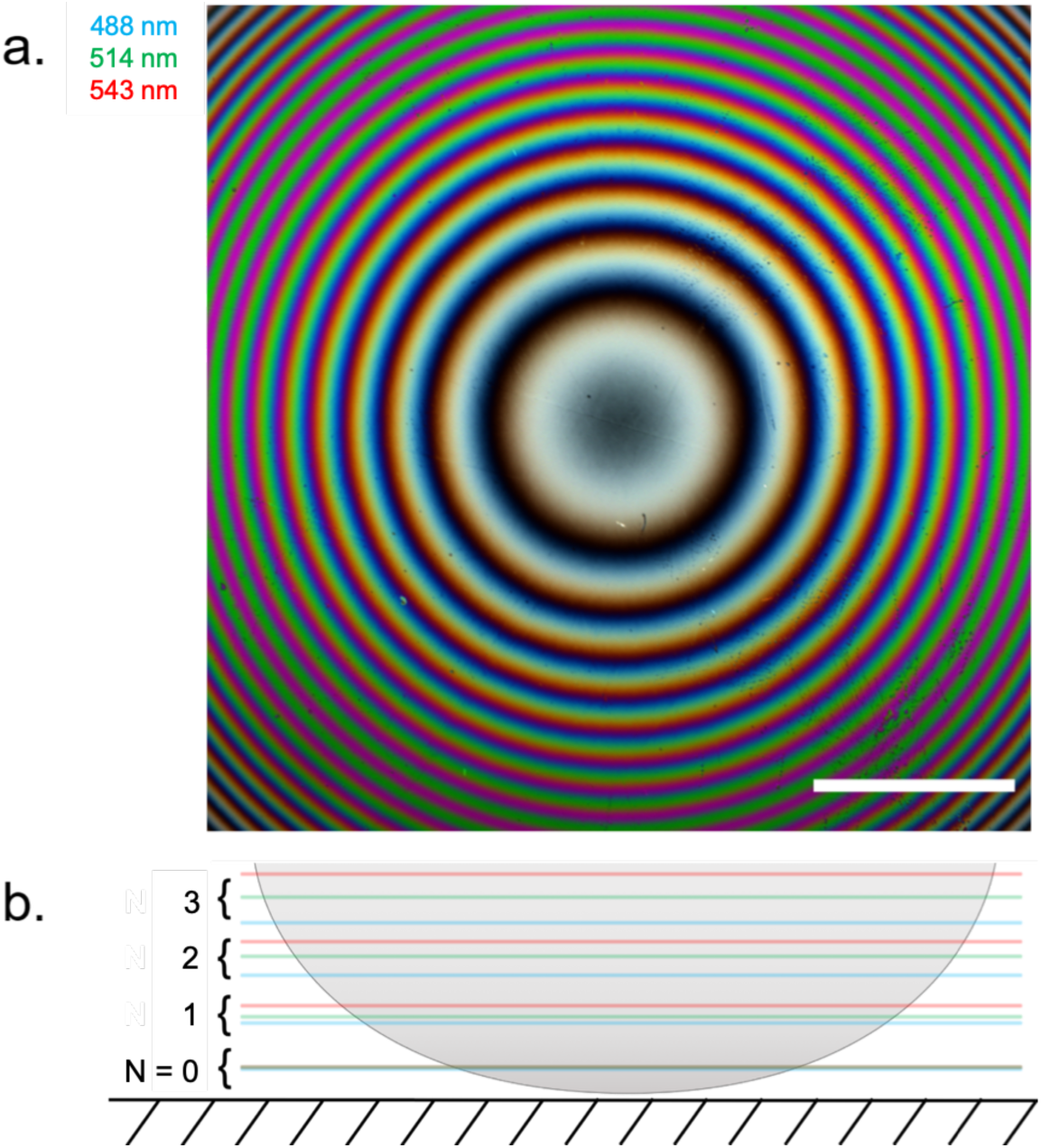
IRM image and schematic diagram of a plano-convex lens specimen. (**a**) A composite IRM image acquired using wavelengths at 488 nm, 514 nm and the 543 nm which are false-coloured blue, green and red. As the concentric fringes propagate away from the coverglass we observed spectral separation of the fringes. Scale bar = 200 µm (**b**) A cross-sectional schematic of the lens specimen showing the colour-ordering of each acquisition wavelength as they propagate axially from the coverglass.

**Figure 2.**
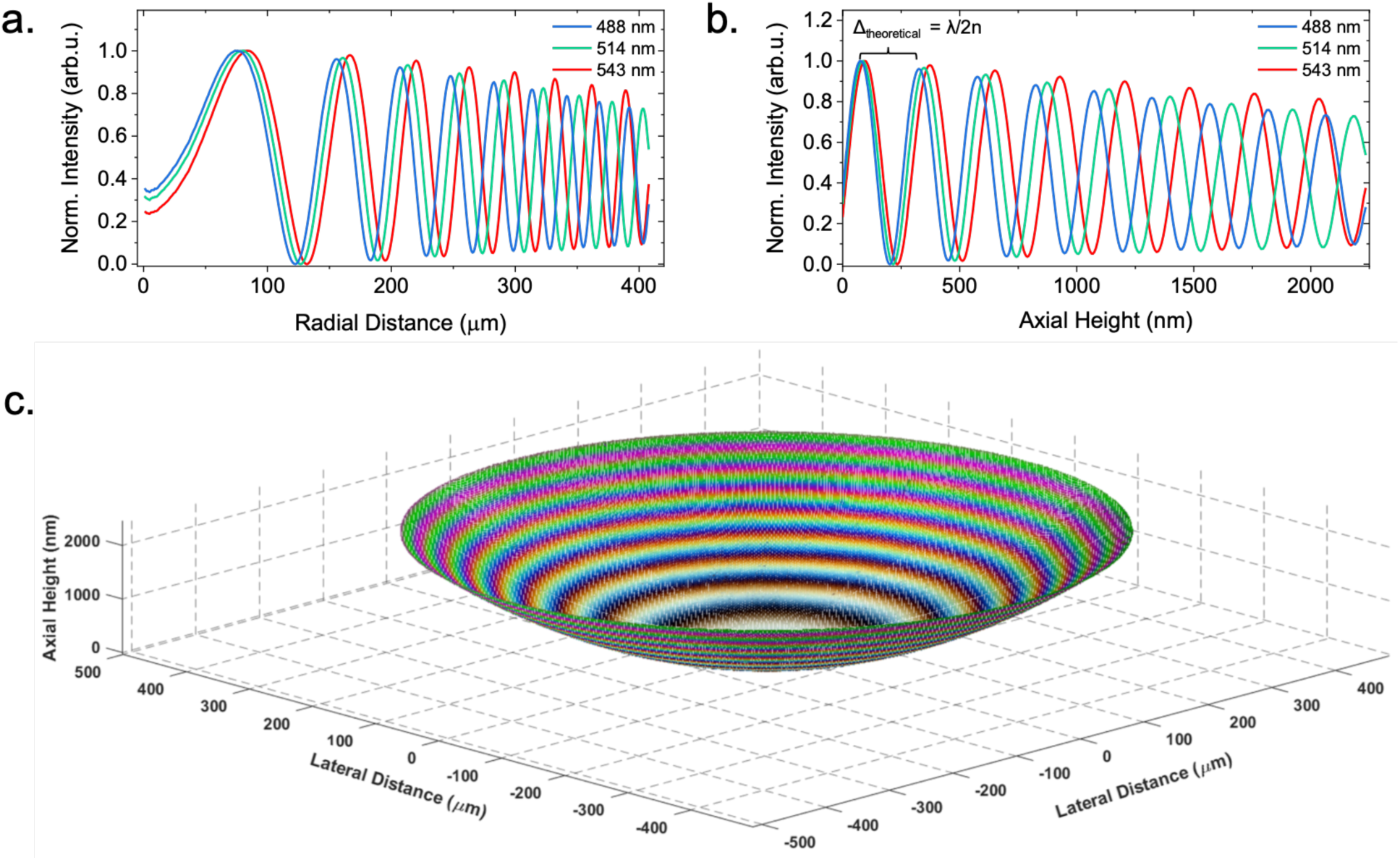
3D reconstruction of a plano-convex lens specimen. (**a**) The radial intensity profile of the interference fringe pattern shown in Figure 1(a). We observed the fringe periodicity decrease as observed in the RGB IRM image. (**b**) Using *a priori* knowledge of the lens specimen, the axial height was calculated and used to plot the intensity of each pixel as a function of height. (**c**) A 3D reconstruction of the lens specimen using the known *x, y* and *z* values and intensity extracted from the 2D IRM image.

### Axial changes along the cell body during gliding motility is independent of AglQ

To demonstrate the benefit of using confocal IRM over widefield IRM we first imaged wild-type *M. xanthus* on a commercial widefield system. We demonstrated that low contrast images are generated in widefield IRM where interference fringes along the cell cannot be resolved clearly. The raw widefield data is presented with a magnified region showing gliding cells (Supplementary Figure 2a) accompanied by the background-corrected data (Supplementary Figure 2b). Interference fringes cannot be clearly resolved in either the raw or corrected data, meaning that a widefield approach is not suitable for studying the changing adhesion profile of gliding bacterial cells.

Using confocal IRM we observed previously undocumented changes in the axial position of *M. xanthus* cells as they glide. Figure 3a shows a single frame from a background-corrected (refer to Supplementary Information) multi-wavelength confocal IRM time-lapse recording of wild-type cells with a magnified view of a representative cell at different time points throughout the time-series (full data set is provided in Supplementary Movie 1). In Figure 3a, clearly resolved interference fringes along the cell body indicate that the cell is not completely attached to the coverglass, but that the cell-to-coverglass distance varies along the body. There is a clear aperiodic change in the fringe pattern over time which indicates changes in the adhesion profile of the cell body during gliding. Interpreting the colour ordering of these fringes, as with the lens specimen, shows that part of the cell body is lifted up from the glass substrate. This observed change in adhesion profile opposes the current theory that gliding cells are firmly adhered along the cell body as they glide. Height changes also show no synchronicity between nearby gliding cells.

**Figure 3.**
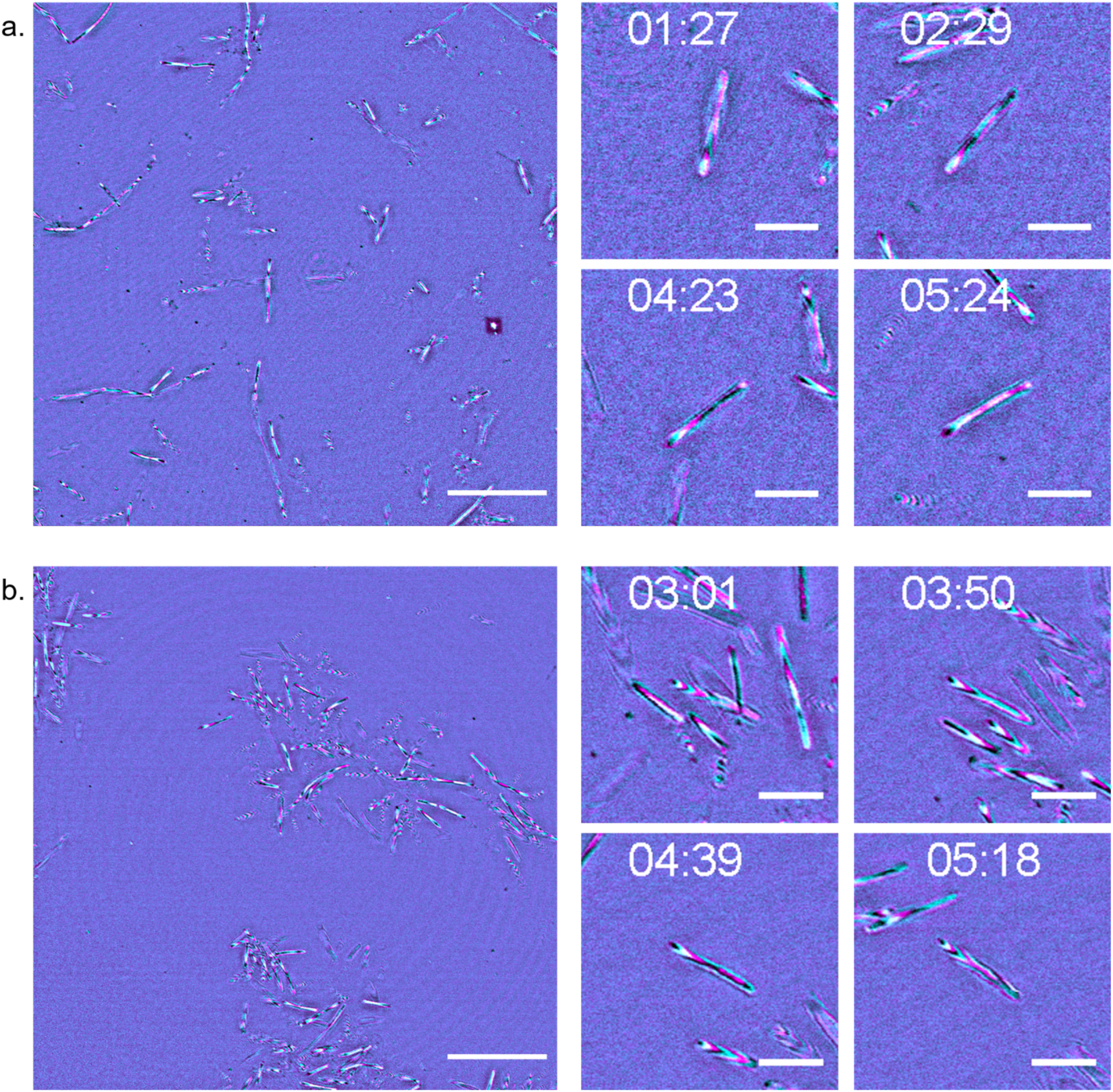
IRM reveals axial movements along the cell body during gliding motility. (**a**) A single frame from a wild type DK1622 gliding specimen with 4 magnified regions of interest (ROI) of a single representative cell over the course of the time-lapse (from t = 1 min 27 s to t = 5 min 24 s). Images were acquired using a multi-wavelength approach, with reflected 488 nm signal false-coloured in cyan and reflected 635 nm signal shown in magenta. As the cell glides across the solid substrate the interference fringe pattern changes as the relative position of the cell to the coverglass fluctuates. (**b**) A single frame of DK1622-Δ*aglQ* with magnified ROI from a single representative gliding cell over the course of the time-lapse (from t = 3 min 1 s to t = 5 min 18 s). Images were acquired using incident light at 488 (cyan) and 635 nm (magenta). DK1622-Δ*aglQ* exhibits the same axial movements as the wild type, demonstrated by the presence of interference fringes which fluctuate as the cell glides. Full time series data for DK1622 and DK1622-Δ*aglQ* are presented in Supplementary Movie 1 and 2, respectively. Scale bar = 20 µm (single frame), 5 µm (ROI).

Figure 3b shows a single frame from a background-corrected multi-wavelength confocal IRM time-lapse recording of a DK1622-Δ*aglQ* strain with a magnified view of a representative gliding cell (full data set is provided in Supplementary Movie 2). The deletion of *aglQ* results in a loss of PMF from the AglRQS complex, which in turn prevents rotation of the MreB cytoskeleton and transit of the FAC along the cell body^23^. Figure 3b shows that the fluctuations in cell topology present in the wild-type are also present in DK1622-Δ*aglQ*. This conserved motility behaviour suggests that axial changes in the cell body during gliding are independent of PMF-driven translocation of FACs along the cell body and are not linked to the proposed membrane protrusions reported in other studies. We then investigated if T4P were responsible for the changes in adhesion profile we report. Deletion of the T4P subunit PilA resulted in cells being unable to adhere to the coverglass and initiate gliding (Supplementary Movies 3 and 4). We reason that force transmission via T4P extension, attachment and retraction could be responsible for the height fluctuations we report.

### Using multi-wavelength IRM for extracting 3D-directionality

Figure 4 illustrates how multi-wavelength IRM data can be assessed qualitatively to understand the geometry of a single gliding *M. xanthus* cell (additional gliding morphologies are presented in Supplementary Figure 3). A representative wild-type gliding cell is presented where multi-colour fringes can be observed along the cell body (Figure 4a). In the IRM image, lifting of the cell body is clearly indicated by an alternating fringe pattern along the body. By interpreting the intensity plot profile along the cell body (Figure 4b) we can extract qualitative topological information about the cell morphology. Figure 4b shows a cyan fringe (λ = 488 nm) followed by a magenta fringe (λ = 635 nm) and based on this we can conclude that the cell body was attached to the coverglass at the leading pole before the basal surface raised to a height of approximately 180 nm. A cartoon diagram approximating the shape of the cell body is shown in Figure 4c. The IRM data indicates a variety of cell orientations and shapes which occur during gliding motility. Examples of additional morphologies are presented in Supplementary Figure 3. These include an undulating topology where the cell is attached to the coverglass at the leading pole while the cell body raises to a height of approximately 180 nm before falling to 90 nm, and again raising to 180 nm at the lagging pole (Supplementary Figure 3a). Another cell motility behaviour is depicted at the point of surface attachment prior to the start of gliding where the leading pole of the cell is attached to the coverglass and the cell body projects upwards into the liquid medium at a sharp incline of approximately 60° relative to the coverglass (Supplementary Figure 3b).

**Figure 4.**
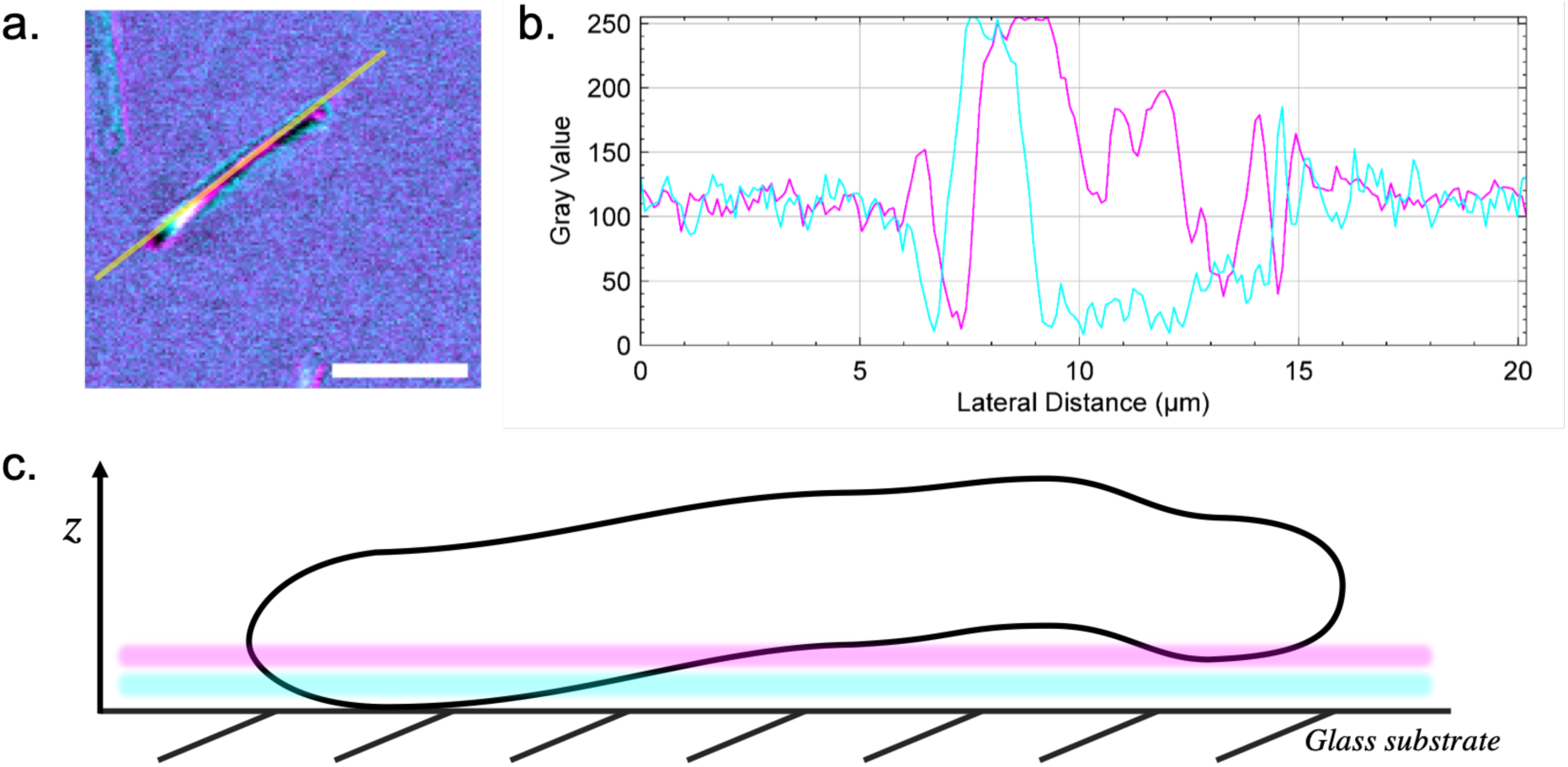
The interference fringe patterns of a gliding cell reveals the axial profile of the cell. (**a**) A representative DK1622 cell is shown from a time series dataset with the location where the intensity profile in (b) was measured. Interference fringes along the cell body can be observed, with reflected 488 nm signal shown in cyan and 635 nm signal in magenta. Scale bar = 5 µm. (**b**) Intensity plot profile from the line through the cell presented in (a). The plot shows the maxima and minima of the interference fringes acquired using both 488 nm (cyan) and 635 nm light (magenta). The spectral separation of the two interference patterns can also be observed. Axial directionality of the cell can be determined by interpreting the colour-ordering of the fringes, where fringes arising from the longer wavelength appear after those from the shorter wavelength when the cell in inclined, and the opposite for declining slopes. The plot was acquired by averaging the signal over a line width of 3 pixels. (**c**) A schematic of the *x, z* profile of the cell shown in (a) according to the colour ordering and intensity profile in (b). The axial position of the coloured fringes from each acquisition wavelength are also shown. The cell is not adhered to the solid surface during gliding and does not maintain a linear cylindrical profile along the length of the cell body. According to theory, regions which intersect the axial position of the first order 488 nm maxima are located 91.7 nm above the substratum, and at 119.3 nm for the first order 635 nm maxima. The schematic is not drawn to scale.

### Using IRM to measure the velocity of gliding cells

We used IRM to determine the mean velocity of motile cells and showed that deletion of *aglQ* decreased the length of time which cells remain adhered to the glass substrate compared to the wild-type (Figure 5a). This implies that AglQ is responsible for maintaining adherence to the coverglass during gliding. The mean velocity of gliding cells was determined by measuring the displacement of the cells over time as they glide, selecting cells with an approximately linear trajectory. We found a 41.2% decrease in mean velocity of DK1622-Δ*aglQ* (mean velocity = 9.26 ± 0.72 µm/min) when compared with the wild-type (mean velocity = 15.76 ± 0.89 µm/min), which concurs with their altered motility phenotype (Figure 5b).

**Figure 5.**
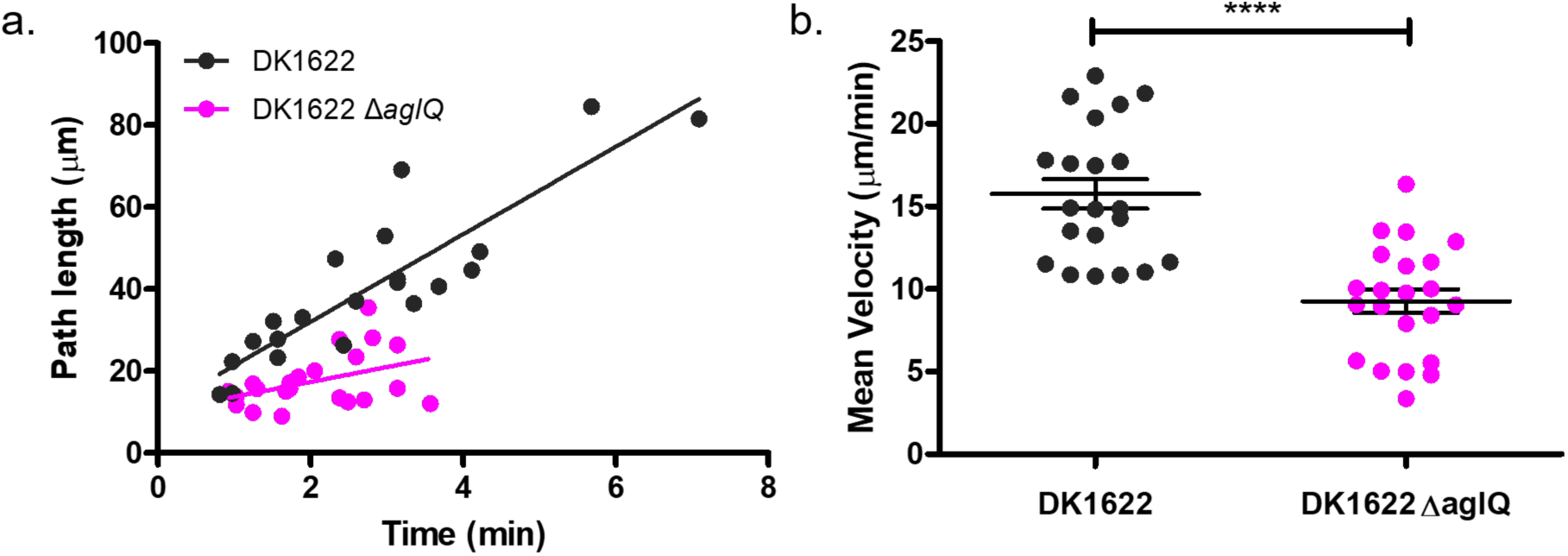
IRM as a method for measuring the velocity of adherent cells. (**a**) The path length of DK1622 and DK1622-Δ*aglQ* are presented over the course of a time series. Overall DK1622 has an increased mean path length when compared to DK1622-Δ*aglQ*, which shows that DK1622-Δ*aglQ* has a lower adhesion profile than the wild type. (**b**) Deletion of *aglQ* results in a decrease in mean velocity of 41.2% from 15.76 ± 0.89 µm/min to 9.26 ± 0.72 µm/min when compared with the wild type. *n*_DK1622_ = 21, *n*_DK1622-ΔaglQ_ = 22, p > 0.0001.

## Discussion

We report aperiodic changes in the adhesion profile of gliding myxobacteria using the label-free technique, IRM. We began with the hypothesis that previous studies had failed to identify any axial behaviours in gliding cells due to the drawbacks with conventional imaging techniques, and that changes in cell height may arise due to the complex mechanisms which govern gliding motility. The data presented here shows new behaviour in gliding myxobacteria which do not fit the current gliding motility models. The behaviours we report suggest that there are additional factors which mediate gliding motility and show the benefit of using IRM to extract 3D information from bacterial specimens by using an easily-implemented microscopy technique.

The current consensus is that gliding cells must be firmly attached to a solid surface to facilitate gliding, however these data show that this is not the case. Our results show that throughout gliding, cells undergo changes in the axial position of their basal surface on the order of 90-180 nm. This new information indicates that the helical rotation and FAC motility models do not fully explain the mechanisms of bacterial gliding. One previous study has used RICM to investigate the adhesion profile of myxobacteria during detachment^17^. However, they did not report any fringes along the cell body, or any aperiodic changes in height in gliding cells. This is likely due to the poor contrast of widefield RICM compared to confocal IRM that prohibits the detection of higher interference fringe orders. It is important to also note that some researchers have claimed that the ability of IRM to detect regions of close contact is questionable due to the inhomogeneous refractive index of the cytosol proximal to the plasma membrane and from self-interference from higher fringe orders within the cell^42,44^. However, these claims are solely based on observations of mammalian cells using widefield IRM. Owing to the lack of vesicular trafficking and much smaller relative size of bacteria compared to mammalian cells, we maintain that confocal IRM remains a valid technique for studying surface adhesion in prokaryotes.

Previous studies which have used conventional optical microscopy methods to image gliding myxobacteria have also failed to observe changes in surface adhesion due to the elongation of the axial point spread function being on the same order as the thickness of a bacterial cell^49,50^. By adopting a confocal approach to IRM for bacterial imaging we overcome the limits of optical sectioning of thin specimens in standard optical microscopy, while simultaneously visualising the adhesion profile of cells. The drawback of using confocal IRM is a decreased temporal resolution when compared with widefield IRM. However, we were still able to track gliding cells sufficiently. While the contrast improvement provided by using a confocal approach over widefield should make image analysis easier^50,51^, the complexity of IRM image data makes image processing and analysis challenging. There are currently no readily available image processing tools that can extract 3D information from a 2D image, such as in IRM. However, with these improved data it may be possible to develop software tools to reconstruct the 3D topography of bacteria using IRM data. The temporal resolution could be improved by using a spinning disk confocal microscope for IRM, however most commercial spinning disk microscopes may not be suitable to incorporate IRM due to hardware limitations^41^. Using confocal IRM also allows us to confirm that the studied cells are in close proximity to the surface of the coverglass and that the gliding behaviours we report are not caused by other factors such as inertia or Brownian motion.

Changes in the relative position of the basal cell membrane above the coverglass could perhaps be explained by proposed distortions in cell wall due to the translocation of FACs along the basal surface of the cell^19^, or because of cell surface unevenness^52^. However, when we compare the axial movements displayed by the wild-type with the Δ*aglQ* mutant we see that this novel gilding behaviour remains. The role of AglQ in gliding motility is central to both the helical rotation and FAC models^22,24,25,53^, and having both wild-type and DK1622-Δ*aglQ* display the same behaviour highlights that more understanding of the mechanisms of gliding motility is required. We then investigated the role of T4P in the changes we observe. Time lapse imaging data was acquired of DK1622-Δ*pilA* and DK1622-Δ*aglQ*, Δ*pilA* mutants to establish if T4P-mediated events altered the adhesion behaviours we observed, but these cells were not able to adhere to the coverglass and did not display an adherence pattern when imaged using IRM. We were therefore unable to confirm if T4P extension, attachment and retraction were responsible for the behaviours we document (Supplementary Movies 3 and 4). We measured the mean velocity of wild-type and DK1622-Δ*aglQ* cells. Using Δ*aglQ* as a control, where we know from previous work that gliding should be impaired^22^, we have shown that height-changing behaviours are independent of the gliding velocity or path length where cells remain associated with the coverglass. To determine the gliding velocity, we measured the average time cells glided along approximately linear trajectories. Routine automated tracking algorithms generally have difficulties in tracking bacterial specimen due to their reliance on blob detection of spherical objects^54–56^. Attempting to isolate and track rod-shaped objects, such as *M. xanthus*, has proven difficult and the addition of interference fringes to rod-shaped objects only adds to the complications of automated cell tracking and analysis. We therefore used a manual tracking method to measure the mean velocity of gliding cells. We concluded that the DK1622-Δ*aglQ* moved on average 30% slower than the wild-type. Also, the wild-type remains adhered to the surface for longer periods of time, yet the dynamic fluctuations in height we report remain.

This work has provided new insights into the 3D motility of bacteria and identified novel motility behaviours in *M. xanthus* which suggest there may be unknown mechanisms which do not agree with the current FAC and helical rotation models. This study does not establish if the novel 3D motility behaviour is only limited to *M. xanthus*, or aperiodic oscillations in height are a behaviour common to other gliding bacteria. We suggest that the fluctuations in height we observe are mediated by T4P and occur due to recoil following firing of pili from the leading pole. In this work we attempted to image T4P mutants to confirm this hypothesis (Supplementary Movies 3 and 4), however these mutants were unable to attach to the glass substrate and therefore unable to glide. Therefore, this study does not resolve the precise mechanism(s) which underpin this behaviour, and further investigation is required to identify factors which play a role in the gliding motility of these organisms. Moving forward it would be interesting to assess the role of T4P in the changing adhesion profile of gliding cells. One way to achieve this would be by investigating the spatiotemporal dynamics of T4P firing in association with the changing adhesion profile of gliding cells using a correlative TIRF-IRM approach. This would allow for simultaneous imaging of the adhesion profile of gliding cells and fluorescently-tagged T4P proximal to the coverglass. The development of an image processing workflow to extract 3D information from confocal IRM images of myxobacteria would also provide users with a method to quantify the aperiodic oscillatory behaviour.

## Materials and Methods

### Bacterial cell culture

*Myxococcus xanthus* cultures (see Table 1) were maintained on double casitone yeast extract (DCYE) medium (20 g/L casein hydrolysate, 2 g/L yeast extract, 8 mM MgSO_4_, 10 mM Tris-HCl, (with 20 g/L agar for solid medium)). For imaging, cells were inoculated at high cell densities in liquid DCYE and grown for 48 hours at 30°C while shaking at 250 rpm. Prior to imaging an 800 μL sample of the exponentially-growing culture was removed from liquid culture and placed in a 35 mm optical-bottom petri dish with a coverslip thickness of 180 µm (cat. no. 80136; ibidi GmbH, Germany) and incubated at 30°C for 20 minutes to allow cells to adhere.

**Table 1.**
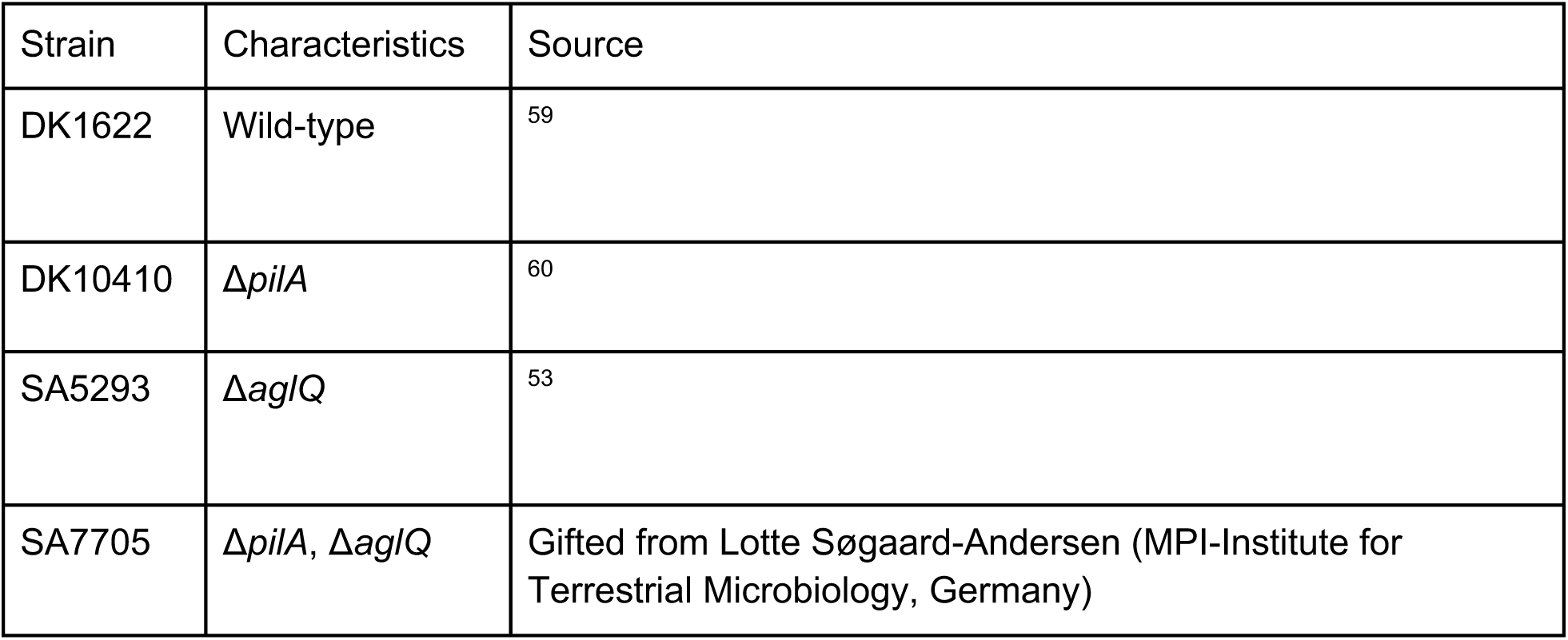
Bacterial strains used in this study.

The refractive index of liquid DCYE medium was measured to be 1.33 using an Abbe Refractometer (Billingham & Stanley Ltd., U.K.).

### Interference Reflection Microscopy

For the characterisation of a model lens specimen, the specimen was placed convex-side down on a 170 µm-thick coverglass measuring 50 × 24 mm which bridged the microscope stage insert. An Olympus IX81 inverted microscope coupled to an Fluoview FV1000 confocal scanning unit (Olympus, Japan) was used to image the lens specimen. The microscope was configured for IRM by replacing the emission dichroic by an 80/20 beam splitter. Images were acquired using a 10x/0.3 N.A. UPlanFl objective lens (Olympus, Japan) and reflection signals were detected using a photomultiplier tube (PMT) for each wavelength, with spectral detection limited to a 10 nm bandwidth over the wavelength of incident light used. A 488 nm line from an Argon laser source (GLG3135; Showa Optronics, Japan) was used for single wavelength acquisition. For multi-wavelength acquisition, 488 nm and 514 nm lines were provided by an Argon laser and 543 nm was provided by a Helium-Neon-Green laser source (GLG3135; Showa Optronics, Japan).

For *M. xanthus* imaging, optical-bottom dishes were placed on the stage of an inverted Olympus IX81 microscope coupled to a Fluoview FV1000 confocal laser scanning unit (Olympus, Japan). Images were acquired using a 60x/1.35 N.A. UPlanSApo Oil objective lens (Olympus, Japan). The microscope was configured for IRM as described above with incident light of 488 nm from an Argon laser and 635 nm obtained from a 635 nm laser diode (GLG3135; Showa Optronics, Japan) for multi-wavelength acquisitions. These wavelengths were selected based on their large spectral separation, meaning that colour-ordering in the observed IRM images was more distinct. For multi-wavelength images both channels were acquired simultaneously with two separate photomultiplier tube detectors.

For widefield IRM specimens were prepared as above and imaged using a Nikon Eclipse-Ti2 inverted microscope (Nikon Instruments, USA) coupled to a Prime 95B sCMOS detector (Teledyne Photometrics, USA). Images were acquired using a 60x/1.4 N.A. PlanApo Oil objective lens (Nikon Instruments, USA). The microscope was configured for IRM by placing an 80/20 beam splitter in the detection path and incident light was sourced from 450 nm and 550 nm LEDs (CoolLED, UK). Multi-wavelength images were acquired sequentially.

### Image processing and analysis

#### Image Correction

A common problem for the analysis of IRM images is the inhomogeneity of brightness across the image that limits the utility of image segmentation tools like thresholding. To correct for changes in image brightness across the field, the moving average of the *k x k* neighbourhood was divided from each pixel using MATLAB 2018b. To rescale the histogram for downstream analysis in FIJI^57^ the image intensity was rescaled by dividing by a factor of 2.

The length of the neighbourhood, *k*, was selected depending on the specimen that was imaged. For the lens specimen, a large *k*-value (*k* = 1000) was chosen to prevent lowering the contrast of the lens signal. For IRM images of *M. xanthus*, where the frequency of the observed interference fringes relative to the pixel density is high, a relatively small *k*-value (*k* = 30) proved suitable. Line intensity profiles from raw and corrected IRM images were checked to verify that the position and intensity succession of the interference fringes were not altered due to the correction method (Supplementary Figures 4, 5 and 6).

#### Lens analysis and reconstruction

In this work, we verified that multi-wavelength IRM can be used to study the change in cell topology during gliding by imaging a lens specimen of known geometry and comparing the results to the theoretical model (refer to Background Theory). The one-dimensional height profile of a lens *z*_*lens*_ follows:

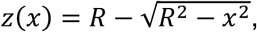

with *R* as the radius of the lens and *x* as the distance to the centre point of the lens that touches the surface/coverslip (*x* = 0, *z* = 0).

Images acquired at multiple wavelengths of the lens specimen were linearly contrast adjusted and cropped to create a square image using FIJI^57^. A composite RGB image was created by merging the channels. The data was imported into MATLAB 2018b, and using the same radial analysis method as presented by Tinning et al., we calculated the axial height of interference fringes from the composite RGB image of the lens specimen^58^. We used a *findpeaks* function to determine the spacing between the experimental intensity maxima of the interference fringes. Each subsequent constructive interference fringe was subtracted from its neighbouring fringe to calculate the experimental spacing.

Lens reconstruction was performed using MATLAB. Firstly, the radial distance for each pixel was extracted from the centre of the RGB IRM image of the lens specimen. By applying *a priori* knowledge of the lens specimen geometry to the radial distance of each pixel (refer to Background Theory), we were able to assign each pixel to an axial height value based on the experimental fringe separation calculated previously. The *x, y* and *z* coordinates along with the image intensities were plotted to create a 3D reconstruction of the RGB IRM image.

## Supporting information

Supplementary movie 2_aglQ

Supplementary movie 3_pilA

Supplementary movie 4_aglQpilA

## Acknowledgements

The authors would like to thank David Whitworth (Aberystwyth University, UK) for helpful discussions and kind gift of the wild-type DK1622 strain used in this work. We would also like to thank Lotte Søgaard-Andersen, Dobromir Szadkowski and Anke Treuner-Lange (Max Planck Institute for Terrestrial Microbiology, Germany) for kind donation of the motility mutants used in this study. Thanks also must be given to the facility staff at the Beatson Advanced Imaging Resource (Beatson Institute, UK), especially to Margaret O’Prey for her technical assistance. This work was supported by the Medical Research Council (MR/K015583/1). LSK is supported by the EPSRC and MRC Centre for Doctoral Training in Optical Medical Imaging (Ref: EP/L016559/1).

## Author Contributions

LMR and LSK acquired the data. LMR, LSK and RS analysed the data. LSK developed the image correction workflow. RS developed the lens analysis scripts. LMR, WBA, PAH and GM designed the study. LMR, LSK, RS, WBA, PAH and GM prepared the manuscript.

## References

1. Skerker, J. M. & Berg, H. C. Direct observation of extension and retraction of type IV pili. Proc. Natl. Acad. Sci. 98, 6901–6904 (2001).

2. Merz, A. J., So, M. & Sheetz, M. P. Pilus retraction powers bacterial twitching motility. Nature 407, 99–102 (2000).

3. McBride, M. J. Bacterial gliding motility: multiple mechanisms for cell movement over surfaces. Annu. Rev. Microbiol. 55, 49–75 (2001).

4. Godwin, S. L., Fletcher, M. & Burchard, R. P. Interference reflection microscopic study of sites of association between gliding bacteria and glass substrata. J. Bacteriol. 171, 4589–4594 (1989).

5. Kasai, T., Hamaguchi, T. & Miyata, M. Gliding Motility of *Mycoplasma mobile* on Uniform Oligosaccharides. J. Bacteriol. 197, 2952–2957 (2015).

6. Chang, L. E., Pate, J. L. & Betzig, R. J. Isolation and characterization of nonspreading mutants of the gliding bacterium *Cytophaga johnsonae*. J. Bacteriol. 159, 26–35 (1984).

7. Hoiczyk, E. Gliding motility in cyanobacteria: observations and possible explanations. Arch. Microbiol. 174, 11–17 (2000).

8. Nan, B. & Zusman, D. R. Novel mechanisms power bacterial gliding motility: Novel mechanisms for bacterial surface motilities. Mol. Microbiol. 101, 186–193 (2016).

9. Nan, B. et al. Myxobacteria gliding motility requires cytoskeleton rotation powered by proton motive force. Proc. Natl. Acad. Sci. 108, 2498–2503 (2011).

10. Nan, B. & Zusman, D. R. Uncovering the Mystery of Gliding Motility in the Myxobacteria. Annu. Rev. Genet. 45, 21–39 (2011).

11. Mignot, T. The elusive engine in *Myxococcus xanthus* gliding motility. Cell. Mol. Life Sci. 64, 2733–2745 (2007).

12. Spormann, A. M. & Kaiser, A. D. Gliding movements in *Myxococcus xanthus*. J. Bacteriol. 177, 5846–5852 (1995).

13. Spormann, A. M. Gliding Motility in Bacteria: Insights from Studies of *Myxococcus xanthus*. Microbiol Mol Biol Rev 63, 621–641 (1999).

14. Muñoz-Dorado, J., Marcos-Torres, F. J., García-Bravo, E., Moraleda-Muñoz, A. & Pérez, J. Myxobacteria: Moving, Killing, Feeding, and Surviving Together. Front. Microbiol. 7, (2016).

15. Zhang, Y., Ducret, A., Shaevitz, J. & Mignot, T. From individual cell motility to collective behaviors: insights from a prokaryote, *Myxococcus xanthus*. FEMS Microbiol. Rev. 36, 149–164 (2012).

16. Mignot, T., Shaevitz, J. W., Hartzell, P. L. & Zusman, D. R. Evidence That Focal Adhesion Complexes Power Bacterial Gliding Motility. Science 315, 853–856 (2007).

17. Faure, L. M. et al. The mechanism of force transmission at bacterial focal adhesion complexes. Nature 539, 530–535 (2016).

18. Pogue, C. B., Zhou, T. & Nan, B. PlpA, a PilZ-like protein, regulates directed motility of the bacterium *Myxococcus xanthus*: A PilZ-like protein regulates *M. xanthus* motility. Mol. Microbiol. 107, 214–228 (2018).

19. Balagam, R. et al. *Myxococcus xanthus* Gliding Motors Are Elastically Coupled to the Substrate as Predicted by the Focal Adhesion Model of Gliding Motility. PLoS Comput. Biol. 10, e1003619 (2014).

20. Islam, S. T. & Mignot, T. The mysterious nature of bacterial surface (gliding) motility: A focal adhesion-based mechanism in *Myxococcus xanthus*. Semin. Cell Dev. Biol. 46, 143–154 (2015).

21. Islam, S. T. et al. Integrin-Like Tethering of Motility Complexes at Bacterial Focal Adhesions. (2018).

22. Sun, M., Wartel, M., Cascales, E., Shaevitz, J. W. & Mignot, T. Motor-driven intracellular transport powers bacterial gliding motility. Proc. Natl. Acad. Sci. 108, 7559–7564 (2011).

23. Nan, B., Mauriello, E. M. F., Sun, I.-H., Wong, A. & Zusman, D. R. A multi-protein complex from *Myxococcus xanthus* required for bacterial gliding motility: Regulation of A-motility in Myxococcus xanthus. Mol. Microbiol. 76, 1539–1554 (2010).

24. Nan, B. et al. Flagella stator homologs function as motors for myxobacterial gliding motility by moving in helical trajectories. Proc. Natl. Acad. Sci. 110, E1508–E1513 (2013).

25. Fu, G. et al. MotAB-like machinery drives the movement of MreB filaments during bacterial gliding motility. Proc. Natl. Acad. Sci. 115, 2484–2489 (2018).

26. Ducret, A., Fleuchot, B., Bergam, P. & Mignot, T. Direct live imaging of cell–cell protein transfer by transient outer membrane fusion in *Myxococcus xanthus*. Elife 2, e00868 (2013).

27. Curtis, A. S. G. The mechanism of adhesion of cells to glass: a study by interference reflection microscopy. J. Cell Biol. 20, 199–215 (1964).

28. Gingell, D. & Todd, I. Interference reflection microscopy. A quantitative theory for image interpretation and its application to cell-substratum separation measurement. Biophys. J. 26, 507–526 (1979).

29. Bereiter-Hahn, J. Quantitative reflection contrast microscopy of living cells. J. Cell Biol. 82, 767–779 (1979).

30. Bailey, J. & Gingell, D. Contacts of chick fibroblasts on glass: results and limitations of quantitative interferometry. J. Cell Sci. 90, 215–224 (1988).

31. Schindl, M. et al. Cell-substrate interactions and locomotion of Dictyostelium wild-type and mutants defective in three cytoskeletal proteins: a study using quantitative reflection interference contrast microscopy. Biophys. J. 68, 1177–1190 (1995).

32. Saunders, R. M. et al. Role of vinculin in regulating focal adhesion turnover. Eur. J. Cell Biol. 85, 487–500 (2006).

33. Sugiyama, N. et al. Label-free characterization of living human induced pluripotent stem cells by subcellular topographic imaging technique using full-field quantitative phase microscopy coupled with interference reflection microscopy. Biomed. Opt. Express 3, 2175 (2012).

34. Saraiva, N. et al. hGAAP promotes cell adhesion and migration via the stimulation of store-operated Ca ^2+^ entry and calpain 2. J. Cell Biol. 202, 699–713 (2013).

35. Amos, L. A. & Amos, W. B. The bending of sliding microtubules imaged by confocal light microscopy and negative stain electron microscopy. J Cell Sci 1991, 95–101 (1991).

36. Simmert, S., Abdosamadi, M. K., Hermsdorf, G. & Schäffer, E. LED-based interference-reflection microscopy combined with optical tweezers for quantitative three-dimensional microtubule imaging. Opt. Express 26, 14499–14513 (2018).

37. Mahamdeh, M., Simmert, S., Luchniak, A., SchäFfer, E. & Howard, J. Label-free high-speed wide-field imaging of single microtubules using interference reflection microscopy. J. Microsc. 272, 60–66 (2018).

38. Fletcher, M. Attachment of *Pseudomonas fluorescens* to glass and influence of electrolytes on bacterium-substratum separation distance. J. Bacteriol. 170, 2027–2030 (1988).

39. Weber, I. Reflection interference contrast microscopy. in Methods in enzymology 361, 34–47 (Elsevier, 2003).

40. Verschueren, H. Interference reflection microscopy in cell biology: methodology and applications. J. Cell Sci. 75, 279–301 (1985).

41. Barr, V. A. & Bunnell, S. C. Interference Reflection Microscopy. Curr. Protoc. Cell Biol. 45, 4.23.1-4.23.19 (2009).

42. Iwanaga, Y., Braun, D. & Fromherz, P. No correlation of focal contacts and close adhesion by comparing GFP-vinculin and fluorescence interference of DiI. Eur. Biophys. J. 30, 17–26 (2001).

43. Mignot, T. & Shaevitz, J. W. Active and passive mechanisms of intracellular transport and localization in bacteria. Curr. Opin. Microbiol. 11, 580–585 (2008).

44. Gingell, D., Todd, I. & Bailey, J. Topography of cell-glass apposition revealed by total internal reflection fluorescence of volume markers. J. Cell Biol. 100, 1334–1338 (1985).

45. Limozin, L. & Sengupta, K. Quantitative Reflection Interference Contrast Microscopy (RICM) in Soft Matter and Cell Adhesion. ChemPhysChem 10, 2752–2768 (2009).

46. Contreras-Naranjo, J. C. & Ugaz, V. M. A nanometre-scale resolution interference-based probe of interfacial phenomena between microscopic objects and surfaces. Nat. Commun. 4, (2013).

47. Talà, L., Fineberg, A., Kukura, P. & Persat, A. *Pseudomonas aeruginosa* orchestrates twitching motility by sequential control of type IV pili movements. Nat. Microbiol. (2019). doi:10.1038/s41564-019-0378-9

48. Raedler, J. & Sackmann, E. On the measurement of weak repulsive and frictional colloidal forces by reflection interference contrast microscopy. Langmuir 8, 848–853 (1992).

49. Wilson, T. & Sheppard, C. Theory and practice of scanning optical microscopy. (Academic Press, 1984).

50. Handbook of Biological Confocal Microscopy. (Springer Science & Business Media, LLC, 2006).

51. Roeder, A. H. K., Cunha, A., Burl, M. C. & Meyerowitz, E. M. A computational image analysis glossary for biologists. Development 139, 3071–3080 (2012).

52. Lünsdorf, H. & Schairer, H. U. Frozen motion of gliding bacteria outlines inherent features of the motility apparatus. Microbiology 147, 939–947 (2001).

53. Jakobczak, B., Keilberg, D., Wuichet, K. & Søgaard-Andersen, L. Contact-and protein transfer-dependent stimulation of assembly of the gliding motility machinery in *Myxococcus xanthus*. PLoS Genet. 11, e1005341 (2015).

54. Chenouard, N. et al. Objective comparison of particle tracking methods. Nat. Methods 11, 281–289 (2014).

55. Tinevez, J.-Y. et al. TrackMate: An open and extensible platform for single-particle tracking. Methods 115, 80–90 (2017).

56. Wolff, C. et al. Multi-view light-sheet imaging and tracking with the MaMuT software reveals the cell lineage of a direct developing arthropod limb. eLife 7, (2018).

57. Schindelin, J. et al. Fiji: an open-source platform for biological-image analysis. Nat. Methods 9, 676–682 (2012).

58. Tinning, P. W., Scrimgeour, R. & McConnell, G. Widefield standing wave microscopy of red blood cell membrane morphology with high temporal resolution. Biomed. Opt. Express 9, 1745 (2018).

59. Kaiser, D. Social gliding is correlated with the presence of pili in *Myxococcus xanthus*. Proc. Natl. Acad. Sci. 76, 5952–5956 (1979).

60. Wu, S. S. & Kaiser, D. Markerless deletions of pil genes in *Myxococcus xanthus* generated by counterselection with the *Bacillus subtilis sacB* gene. J. Bacteriol. 178, 5817–5821 (1996).

