## Supplementary Figures 1-6 for "Three-dimensional observations of an aperiodic oscillatory gliding motility behaviour in *Myxococcus xanthus* using confocal interference reflection microscopy"

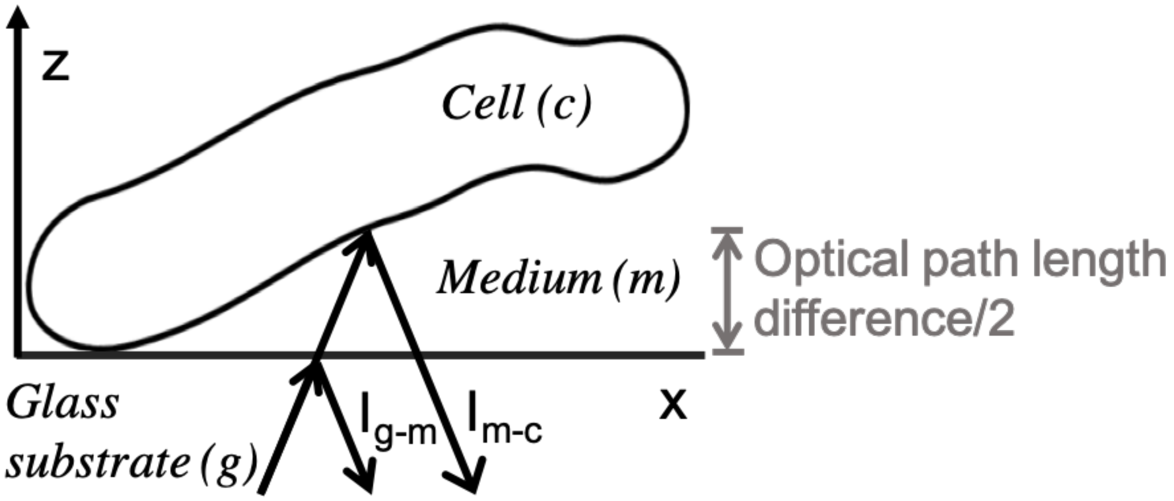


### Supplementary Figures

Supplementary Figure 1. ***Schematic of the IRM model.*** The model system consists of the coverglass, the cell medium and the cell body. Reflections occur at both the coverglass-to-medium and medium-to-cell interfaces. Propagation of direction of the incident and reflected light is perpendicular to the coverglass. The two reflected light beams interfere (de-)constructively, dependent on the optical path length difference.

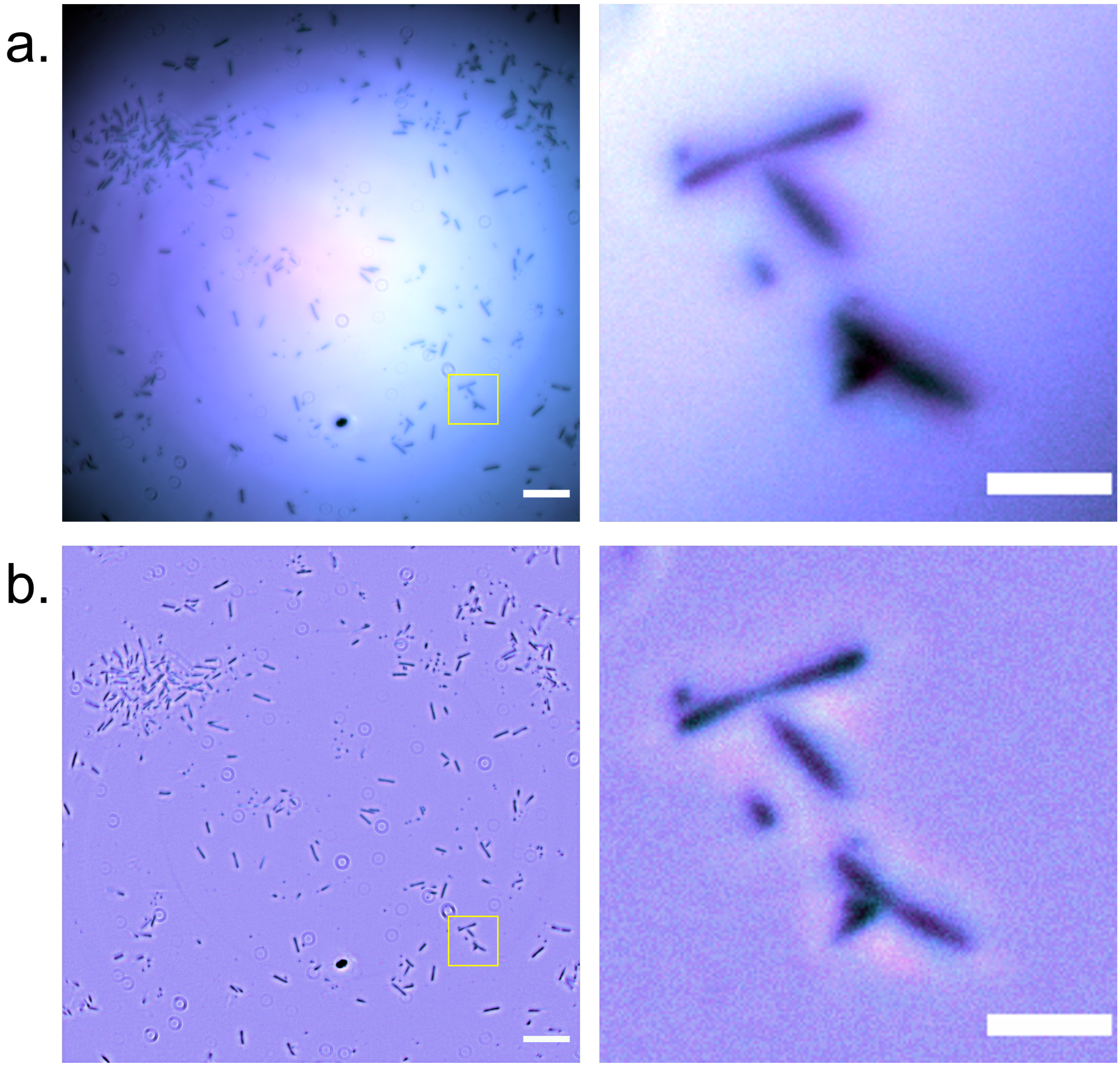


Supplementary Figure 2. ***Widefield IRM does not provide sufficient contrast to reveal sub-diffraction limited changes in the adhesion profile of gliding bacteria.*** Unprocessed **(a)** and background corrected ($k$ = 30) **(b)** multi-wavelength widefield IRM composite image of *M. xanthus* (450 nm = cyan, 550 nm = magenta). Magnified ROI of gliding cells shows low contrast between fringe order when images are acquired using widefield modality. Full field scale bars = 20 μm, ROI scale bars = 5 μm.

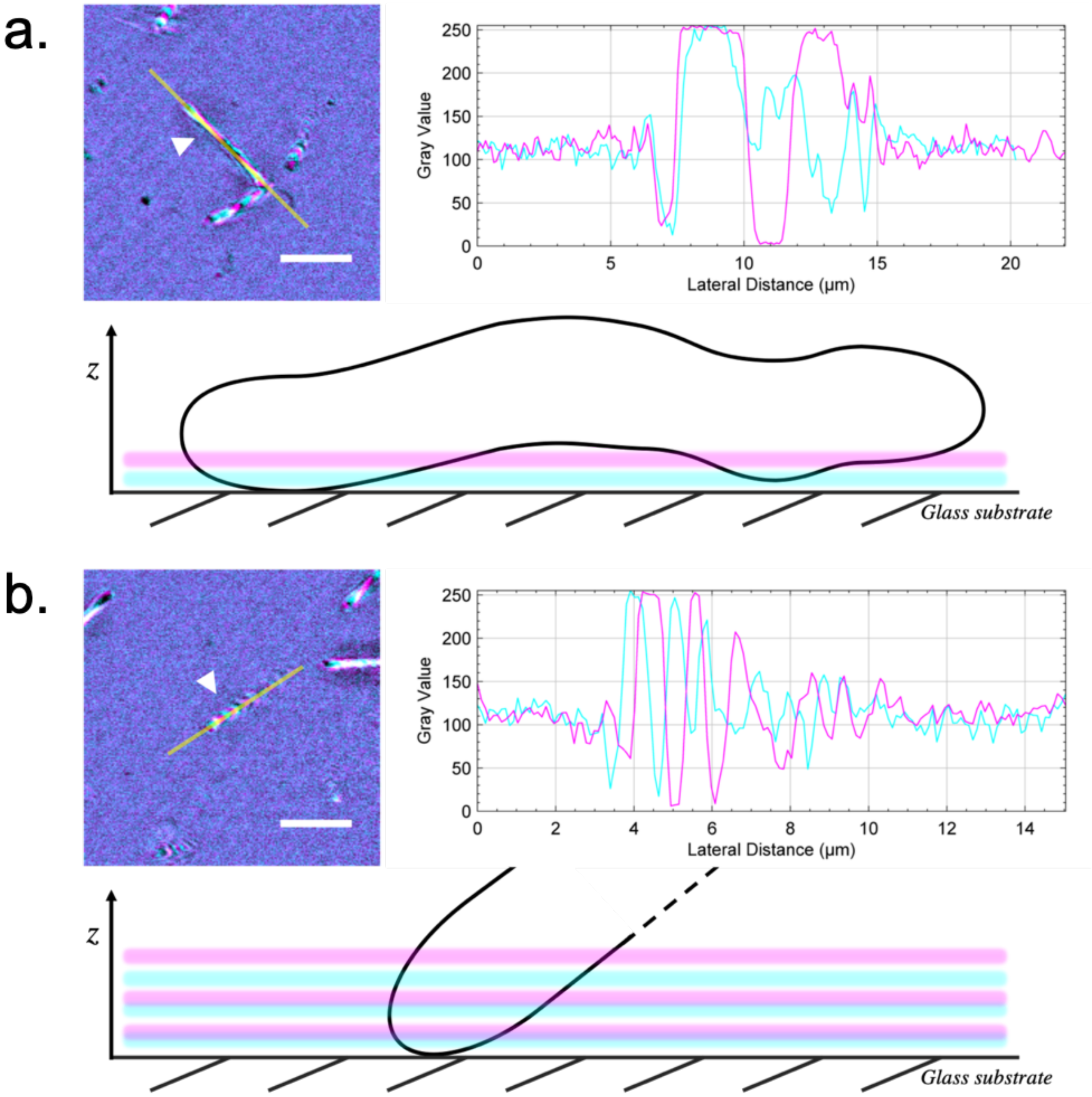


Supplementary Figure 3. ***Additional motility behaviours displayed by Myxococcus xanthus***. (**a**) corrected IRM image of a wild type gliding cell with accompanying intensity plot profile (line width = 3 px (0.3 µm)). An $x$, $z$ schematic is presented which illustrates an undulating motion of the basal surface of the cell during gliding. (**b**) a corrected IRM image of a wild type cell with accompanying intensity plot profile (line width – 3 pixels). The schematic shows the cell attached to the substrate at one pole and projecting upwards into the surrounding medium. Reflected 488 nm light is false-coloured in cyan, and 635 nm light in magenta. Scale bar = 5 $\mu$m.

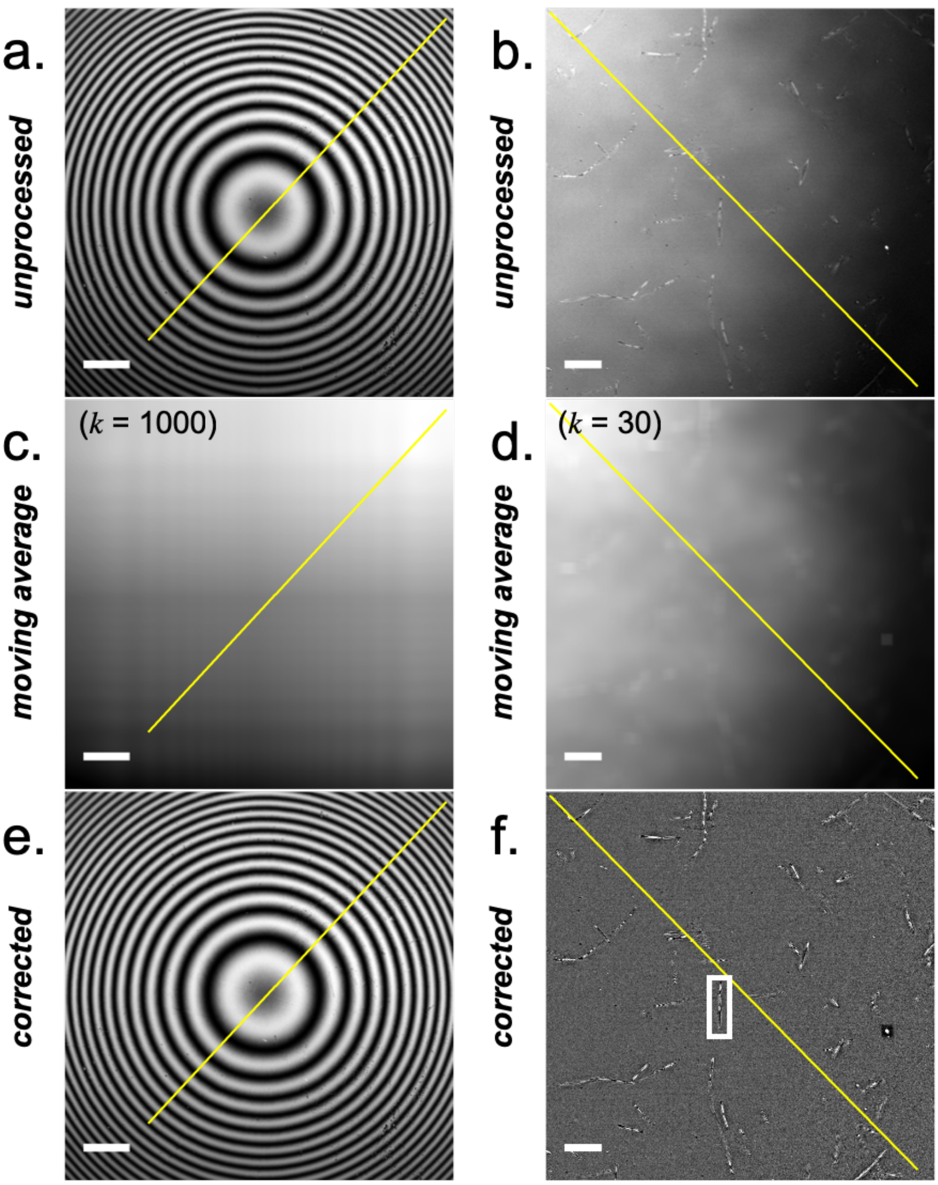


Supplementary Figure 4. ***Image correction workflow for IRM data***. (**a**) Linearly contrast-adjusted IRM image of a 6 mm diameter lens specimen (focal length = 72 mm) acquired using light at 488 nm. Scale bar = 100 μm. (**b**) Linearly contrast adjusted IRM image of *M. xanthus* with inhomogeneous background intensity. Scale bar = 10 μm. (**c**) Calculated moving average of (a) when $k$ = 1000. The calculated moving average accounts for the inhomogeneous background which is subtracted to correct the raw image. (**d**) Calculated moving average of (b) when $k$ = 30. (**e**) Corrected IRM image resulting from (a). (**f**) Corrected IRM image resulting from (b). The ROI selects a single cell where the intensity profile was measured in Supp. Figure 4. Selected line plots are shown in each panel where intensity profiles were measured in Supp. Figure 3.

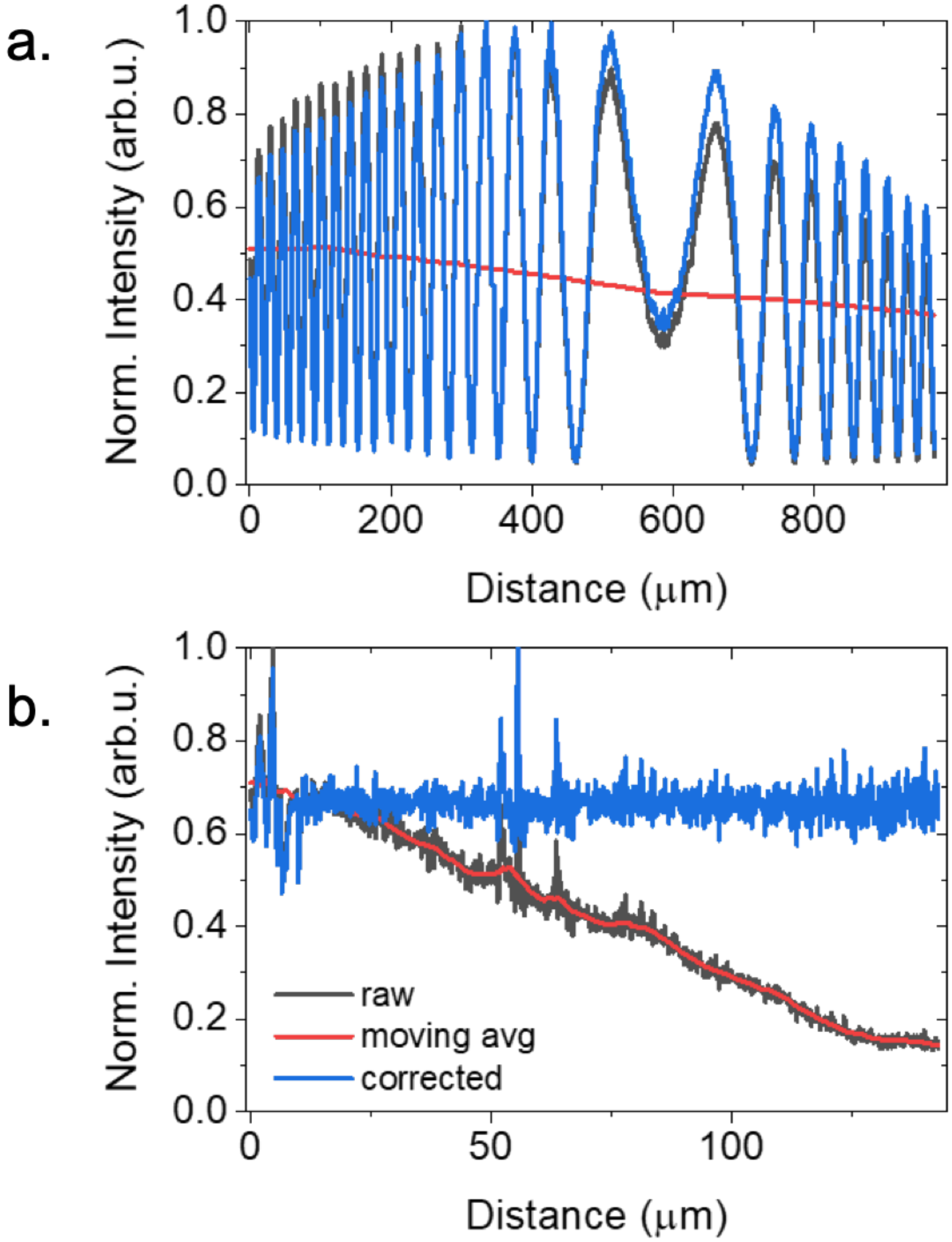


Supplementary Figure 5. ***Image correction does not alter the position of fringe maxima and minima***. (**a**) Selected line intensity profiles from Supp. Figure 1 (a, c and e) are shown. Following correction of the raw images using the calculated moving average, there was no change in the peak position of the intensity maxima and minima. (**b**) Selected line intensity profiles from Supp. Figure 1 (b, d and f) are shown. Correction resulted in homogeneity of illumination across the field. Line intensity profiles were averaged using a line width of (**a**) 5 px (4.2 μm) and (**b**) 5 px (0.5 μm).

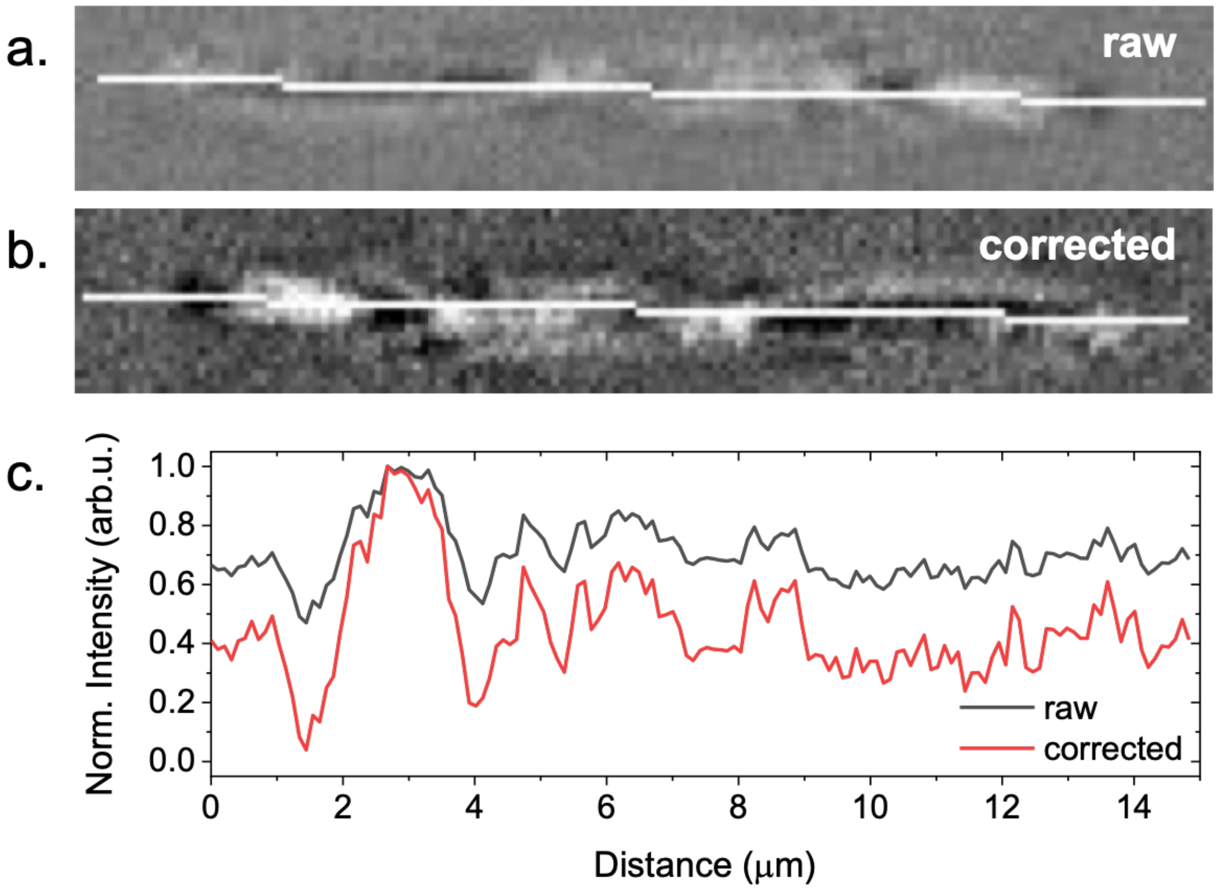


Supplementary Figure 6. ***Image correction does not alter the position of intensity peaks***. (**a**) Unprocessed ROI from Supp. Figure 1(f) with a line profile shown for reference. (**b**) Corrected ROI shown in (a) ($k$ = 30). (**c**) Selected line intensity profiles of (a) and (b) show that there are no changes to the position of intensity values along the length of gliding cells. Line intensity profiles were averaged using a line width of 5 px (0.5 μm).

Supplementary Movie 1. **IRM time series of gliding DK1622 cells**.

Supplementary Movie 2. **IRM time series of gliding DK1622-Δ*aglQ* cells**.

Supplementary Movie 3. **IRM time series of gliding DK1622-Δ*pilA* cells**.

Supplementary Movie 4. **IRM time series of gliding DK1622-Δ*aglQ*, Δ*pilA* cells**.
